# SARS-CoV-2 escapes CD8 T cell surveillance via mutations in MHC-I restricted epitopes

**DOI:** 10.1101/2020.12.18.423507

**Authors:** Benedikt Agerer, Maximilian Koblischke, Venugopal Gudipati, Mark Smyth, Alexandra Popa, Jakob-Wendelin Genger, Lukas Endler, David M. Florian, Vanessa Mühlgrabner, Alexander Lercher, Pia Gattinger, Ricard Torralba-Gombau, Thomas Penz, Ingrid Fae, Sabine Wenda, Marianna Traungott, Gernot Walder, Gottfried Fischer, Wolfgang Hoepler, Erich Pawelka, Alexander Zoufaly, Rudolf Valenta, Christoph Bock, Johannes B. Huppa, Judith H. Aberle, Andreas Bergthaler

**Affiliations:** CeMM Research Center for Molecular Medicine of the Austrian Academy of Sciences, Vienna, Austria; Center for Virology, Medical University of Vienna, Vienna, Austria; Institute for Hygiene and Applied Immunology, Center for Pathophysiology, Infectiology and Immunology, Medical University of Vienna, Vienna, Austria; Institute of Pathophysiology and Allergy Research, Center for Pathophysiology, Infectiology and Immunology, Medical University of Vienna, Vienna, Austria; Department of Blood Group Serology and Transfusion Medicine, Medical University of Vienna, Vienna, Austria; Department of Medicine IV, Kaiser Franz Josef Hospital, Vienna, Austria; Division of Hygiene and Medical Microbiology, Medical University of Innsbruck, Innsbruck, Austria; Department of Laboratory Medicine, Medical University of Vienna, 1090 Vienna, Austria

## Abstract

CD8+ T cell immunity to SARS-CoV-2 has been implicated in COVID-19 severity and virus control, though direct evidence has been lacking so far. Here, we identified non-synonymous mutations in MHC-I restricted CD8+ T cell epitopes after deep sequencing of 747 SARS-CoV- 2 virus isolates. Mutant peptides exhibited diminished or abrogated MHC-I binding, which was associated with a loss of recognition and functional responses by CD8+ T cells isolated from HLA-matched COVID-19 patients. Our findings highlight the capacity of SARS-CoV-2 to subvert CD8+ T cell surveillance through escape mutations in MHCI-restricted viral epitopes. This provides evolutionary evidence for CD8+ T cell immunity controlling SARS-CoV-2 with consequences for COVID-19 vaccine design.

---

SARS-CoV-2 infection elicits broad activation of the innate and adaptive arm of immunity (Le Bert et al., 2020; Mathew et al., 2020; Vabret et al., 2020; Zhang et al., 2020a). Cytotoxic CD8+ T-Lymphocyte (CTL) responses have been described in great detail in SARS-CoV-2 – infected patients (Le Bert et al., 2020; Braun et al., 2020; Dan et al., 2020; Grifoni et al., 2020a; Mathew et al., 2020; Schulien et al., 2020; Sekine et al., 2020). Numerous human leucocyte antigen (HLA)-restricted CTL epitopes have been characterized for SARS-CoV-2 (Le Bert et al., 2020; Grifoni et al., 2020b; Nelde et al., 2020; Poran et al., 2020; Shomuradova et al., 2020), but whether and how CTLs contribute to protective immunity remains unclear.

CTLs kill infected cells upon recognition of viral peptides as they are displayed on the cell surface in the context of the products of the class I major histocompatibility complex (MHC-I). CTLs play an essential role in conferring immune memory and protection against viral pathogens (Goulder and Watkins, 2008; Schmitz et al., 1999; Thimme et al., 2003). Compelling evolutionary evidence for CTL-mediated control of RNA viruses like HIV and HCV is provided by mutations occurring in viral epitopes which directly interfere with MHC-I restricted T cell antigen recognition and killing by CTLs (Cox et al., 2005; Deng et al., 2015; Goulder et al., 2001; Pircher et al., 1990). The extent to which SARS-CoV-2 mutations may upend the presentation of virus-derived peptides via MHC-I remains to be identified.

To assess the impact of CTLs on the control of SARS-CoV-2, we performed deep viral genome sequencing (> 20.000X coverage (Fig. S1A) and bioinformatic analysis on 747 SARS-CoV-2 patient samples (Popa et al., 2020). We focused on 27 CTL epitopes, which were experimentally shown to be presented by the common subtype HLA-A*02:01 (allele frequency 0.29 in Austria) as well as by the minor subtype HLA-B*40:01 (allele frequency 0.03-0.05 in Austria) (Gonzalez-Galarza et al., 2019). There, we detected 197 non-synonymous mutations present at frequencies of ≥ 0.02 in 233 samples (Fig. 1A-B) (Le Bert et al., 2020; Grifoni et al., 2020b; Poran et al., 2020; Shomuradova et al., 2020). Of these 197 variants, 33 were found at frequencies between 0.1 to 0.5. Notably, 9 variants were fixed and found in 53 different patient samples (Table S1). Due to overlaps in some epitopes, these 197 mutations result in 207 different epitope variants. 27 of these mutations were localized to anchor residues, and 14 mutations affected auxiliary residues, which are both integral to MHC-I peptide loading (Fig. S1B). Prediction of the binding strength of the wildtype and mutant peptides to HLA-A*02:01 and HLA-B*40:01 via netMHCpan v4.1 (Reynisson et al., 2020) revealed weaker peptide binding to MHC-I, as indicated by an increase of netMHCpan ranks (Fig. 1A and S1C-F). For many of the investigated CTL epitopes, we detected multiple variants that independently emerged in different SARS-CoV-2 infected individuals (Fig. 1A). To corroborate these findings from low-frequent mutations in our deep sequencing dataset, we analyzed fixed mutations in >145.000 available global SARS-CoV-2 sequences from the public database GISAID (Elbe and Buckland-Merrett, 2017). Mutations were observed in 0.0000689 - 7.336% epitope sequences (mean = 0.005106) (Fig. 1C). We found 10 to 11717 viral genome sequences with a non-synonymous mutation for each of the investigated 27 CTL epitopes (mean = 807.05). Importantly, we found fixed variants in GISAID that were also identified in our low-frequency analysis, highlighting the relevance of individual low-frequency mutations (Fig. 1D, S1G–S1I). An alanine to valine mutation within the reported LALLLLDRL epitope (underlined) at nucleotide position 28932 was found in 7.32% (11693/159720) of the analyzed sequences (Schulien et al., 2020). First detected in June 2020, this variant is now (12.11.2020) not only one of the mutations defining the 20A.EU1 subclade (Fig. S1J-K) (Hodcroft et al., 2020), but it has also been identified in other virus isolates (Fig. S1J). Importantly, this mutation had already been detected at low frequency in 7 of our samples collected between March and April 2020, again suggesting independent emergence of the same mutation in multiple individuals (Fig. S1I).

**Fig. 1.**
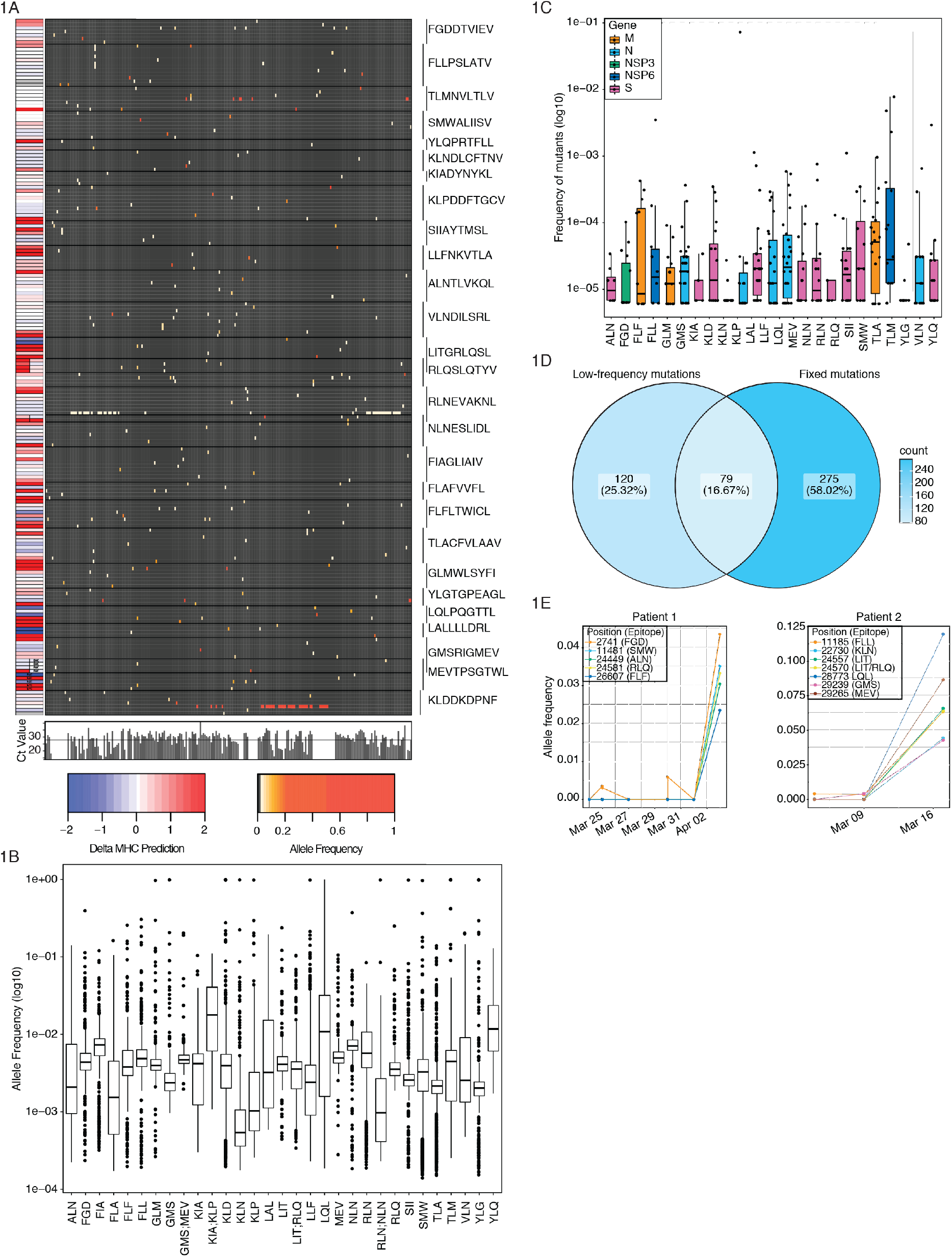
Non-synonymous mutations are detected in SARS-CoV-2 CTL epitopes. **A)** Allele frequency of low-frequency mutations detected in 27 CTL epitopes. Epitopes are indicated on the right. The heatmap to the left indicates change in binding ranks predicted by netMHC-pan 4.1(Reynisson et al., 2020). Barplots below the heatmap indicate viral loads as Ct values. **B)** Allele frequency of mutations in specified epitopes. Regions present in two epitopes are depicted separately. **C)** Frequency of global fixed mutations in CTL epitopes. **D)** Venn diagram depicting overlap between global fixed mutations and low frequency variants **E)** Mutations in CTL epitopes arise late in infection. Mutation frequency over time of two patients which were longitudinally sampled. Shown are variants that lead to non-synonymous mutations in CTL epitopes.

To examine the time dynamics of low-frequency epitope mutations in patients, we utilized serially-sampled viral genomes from COVID-19 patients. We observed that mutations in viral epitopes arising later in infection (Fig. 1F) hinting at CTL-mediated selection pressure mounted over time. Based on our analysis of low-frequency and fixed mutations, we selected 11 wildtype and 17 corresponding mutant peptides with predicted decreased HLA-binding strength for further biophysical and functional analysis (Table S2).

We next employed cell-free differential scanning fluorimetry as a means to compare the capacity of SARS-CoV-2-derived peptides and their non-synonymous mutants to stabilize the structure of recombinant HLA-A*02:01 or HLA-B*40:01 proteins as a direct indicator of MHC-I-peptide binding (Fig. 2A-E, S2A-I) (Blaha et al., 2019). For 9 of the 11 wildtype peptides we observed binding to HLA-A*02:01 or HLA-B*40:01, suggesting that these peptides could potentially be presented by the respective HLA allele. As shown in Fig. S2A, the minima of the curve specify the melting temperature (T_m_) of the HLA-peptide complexes. T_m_ values well above 37°C indicate strong peptide binding to MHC-I at physiological temperatures, whereas values around 37°C correlate only with weak and below 36°C with absent binding. In total 11 analyzed mutants exhibited decreased stabilizing capacity towards MHC-I (Fig. 2A-E, S2D–S2H). The HLA-B*40:01 – restricted MEVTPSGTWL peptide featured specific binding to recombinant HLA-B*40:01 but not to HLA-A*02:01 (Le Bert et al., 2020) (Fig. 2C, S2I). An example for a weak binder is the mutant variant YFQPRTFLL (instead of YLQPRTFLL), whereas LFFNKVTLA (instead of LLFNKVTLA) represents a non-binder for HLA-A*02:01 (Fig. 2A, D). Of note, we did not observe binding of neither wildtype nor mutant peptides for the predicted CTL epitope LALLLLDRL (Fig. S2J).

**Fig. 2.**
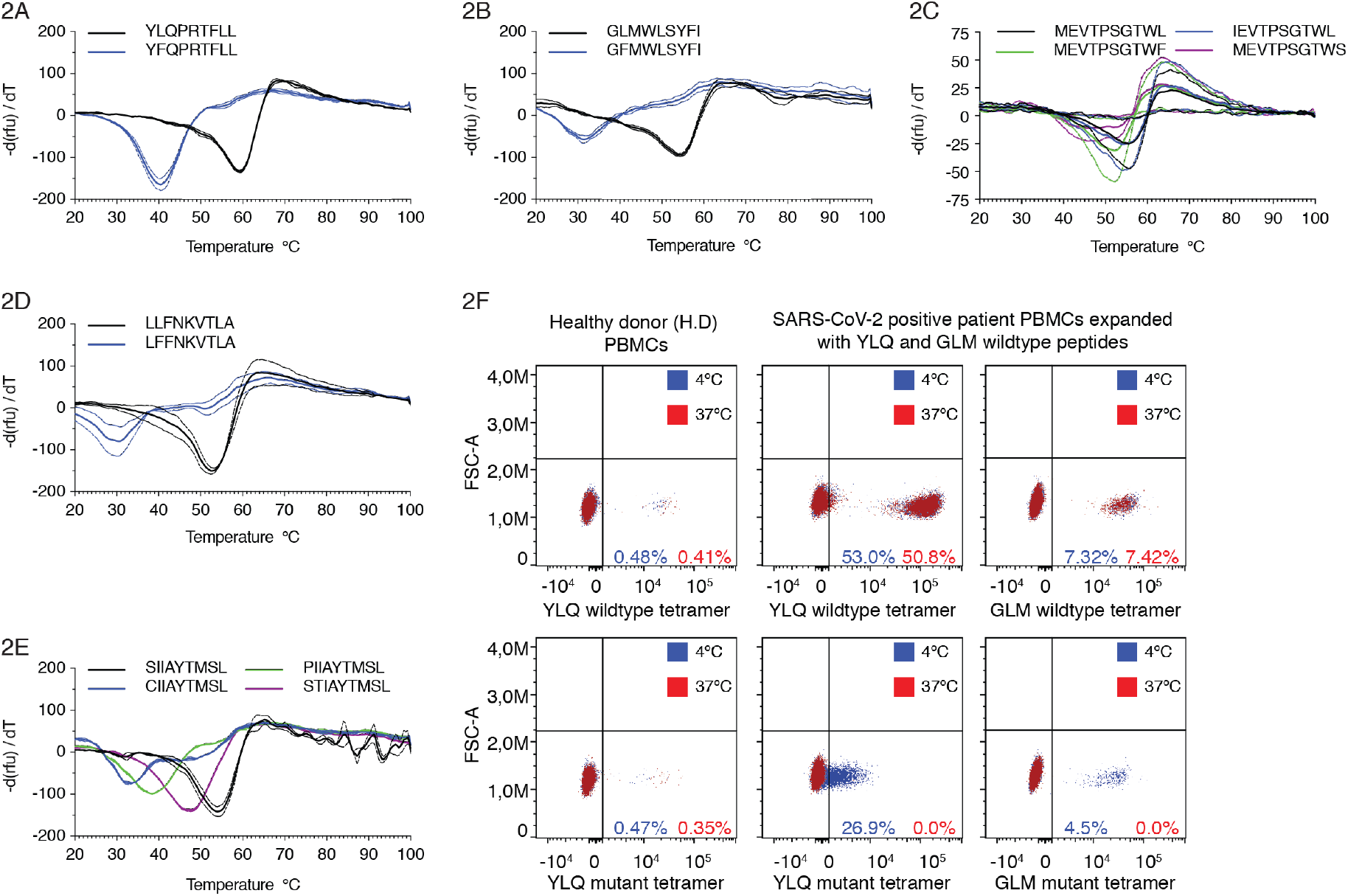
Epitope variants lead to diminished MHC-1 binding. **A-E**) Decreased thermostability of mutant peptide MHC-I complexes. Negative first derivative of relative fluorescence units (rfu) plotted against increasing temperatures. Curves for wildtype peptides are black, mutated peptides are colored. The minimum point of the curves represents the melting temperature of peptide-MHC-I complexes. **F)** Tetramers featuring mutated peptides are instable at 37°C. FACS plots showing staining of in vitro expanded PBMCs stained with tetramers containing wildtype (top) or mutant (bottom) peptides incubated at 4°C (blue) or 37°C (red).

To further corroborate these results, we generated peptide-loaded HLA-A*02:01 and HLA- B*40:01 I tetramers presenting wildtype and mutant peptides as a means to identify cognate CD8+ T-cells from expanded PBMCs of HLA-matched COVID-19 patients (Fig. 3A). As shown in Fig. 2F, tetramers loaded with mutant peptides stain cognate T cells in a TCR-dependent fashion when kept at 4°C. However, when tetramers were incubated at 37°C prior to their use, T cell staining was abrogated, most likely due to peptide loss and structural disintegration of MHC-I. Taken together, these results imply that mutations found in SARS-CoV-2 isolates promote immune escape from HLA-dependent recognition by CTLs.

**Fig. 3.**
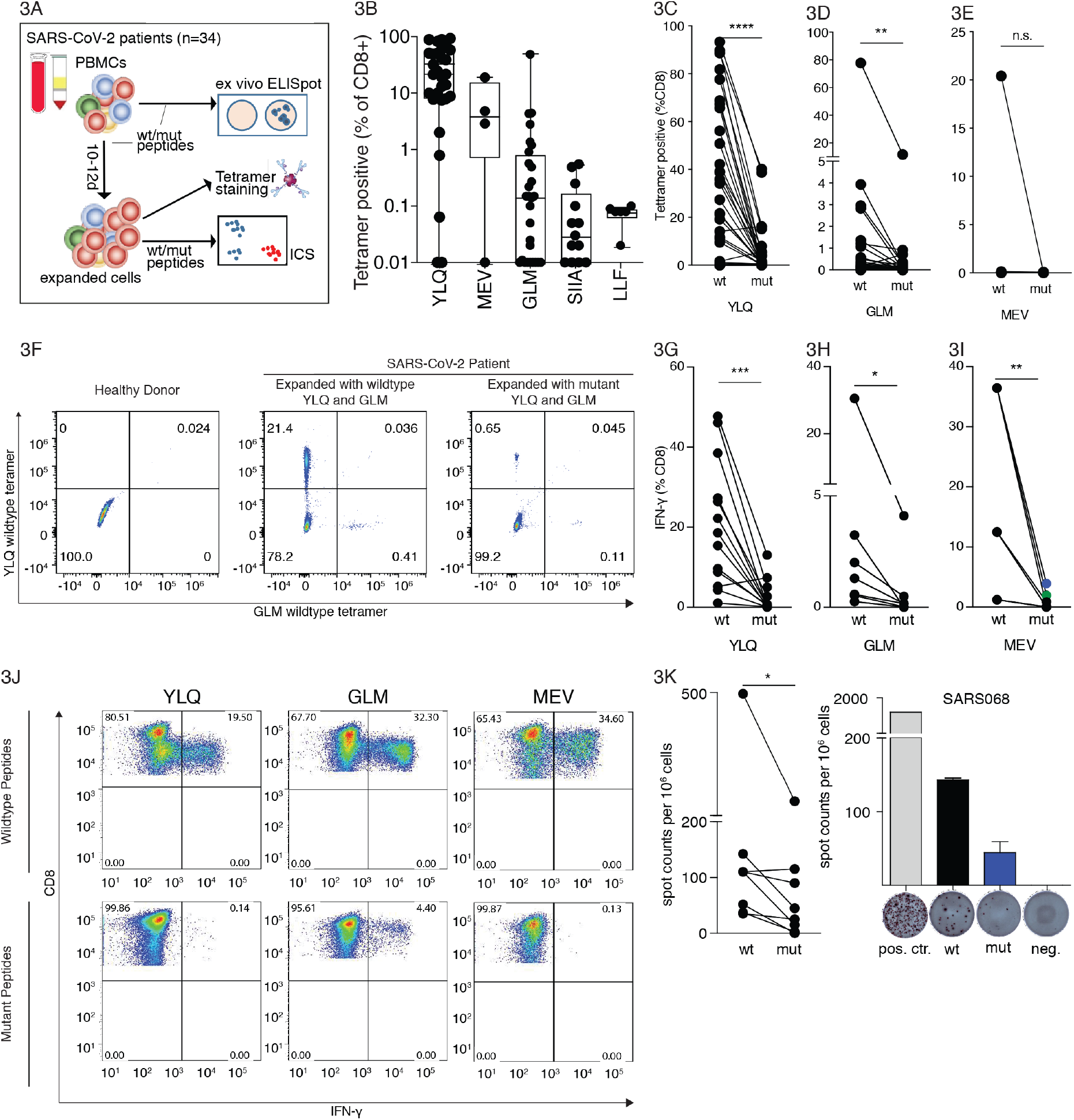
SARS-CoV-2 epitope mutations lead to escape from CTL responses. **A)** Experimental overview. **B)** CTL responses against wildtype epitopes. PBMCs were isolated from HLA- A*02:01 or HLA-B*40:01 positive SARS-CoV-2 patients, expanded 10-12 days with indicated peptides and stained with wildtype tetramers. **C-E)** T-cells expanded with mutant peptides do not give rise to wildtype peptide-specific PBMCs. PBMCs were isolated as in B), stimulated with wildtype or mutant peptides and stained with tetramers containing the wildtype peptide. **F)** Representative FACS plots for C-E. **G-I)** Impact of mutations on CTL cell response. PBMCs expanded with wildtype or mutant peptides as indicated, were analyzed for IFNγ- production via intracellular cytokine staining after restimulation with each peptide (n = 3-13 patients per epitope). **J)** Representative FACS plots for D-F. **K)** *Ex vivo* IFNγ ELISpot assays from PBMCs stimulated with the YLQ peptide or the corresponding mutant. (N=6) or the MEV peptide and corresponding mutant (N=1), shown are results from one repesentative patient for stimulation with the YLQ peptide. Significance is indicated as \**P*<0.05, \*\**P*<0.01, \*\*\**P*<0.001, \*\*\*\**P*<0.0001, tested by Wilcoxon matched-pairs signed rank test

Next, we investigated peptide-specific CD8+ T-cell responses in PBMCs isolated from HLA- A*02:01 or HLA-B*40:01 positive COVID-19 patients (Fig. 3A, Table S3, Table S4). To this end HLA-matched PBMCs from COVID-19 patients were stimulated with peptides and cultured for 10-12 days followed by tetramer staining (Fig. 3B). This allowed us to confirm the wildtype peptides as bona-fide T-cell epitopes in SARS-CoV-2. We further corroborated virus-specific CTL responses by intracellular cytokine staining for IFNγ after peptide-mediated restimulation (Fig. S3A, B). To investigate the extent to which identified mutations in viral epitopes affected T cell proliferation, we stained expanded PBMCs with wildtype peptide-loaded HLA tetramers. We found fewer tetramer-positive CTLs in PBMCs expanded in the presence of mutant peptides, indicating that stimulation with mutant peptides significantly reduces immunogenicity (Fig. 3C-F). Consistent with the tetramer data, intracellular cytokine staining (ICS) for IFNγ revealed significantly diminished CTL responses to mutant peptide as compared to the corresponding wildtype peptides (Fig. 3G-J). We observed markedly diminished CTL responses to several mutant peptides both presented in the context of HLA- A:02*01 and HLA-B:40*01. This was further supported by peptide titration experiments involving wildtype peptides and their mutant counterparts for T-cell stimulation (Fig. S3C). To complement intracellular cytokine measurements, we also carried out *ex vivo* ELISpot assays after stimulating PBMCs with wildtype or mutant peptides without extended expansion. These experiments confirmed that stimulation with mutant peptides resulted in significantly fewer cytokine-producing cells than with wildtype peptides (Fig. 3K, S3D).

Taken together, our data indicate that SARS-CoV-2 may evade CTL surveillance through mutations in viral epitopes which lead to reduced peptide MHC-I binding. Deep SARS-CoV- 2 genome sequencing results afford a valuable additional perspective that complements insights gained from numerous studies on SARS-CoV-2-specific T cell responses (Altmann and Boyton, 2020). Our data does not rule out that altered residues facing the cognate T cell receptor may give rise to the emergence of CTL neoepitopes.

Viruses employ numerous strategies to evade CD8+ T cell immune responses (Alcami and Koszinowski, 2000; Goulder and Watkins, 2004; Hansen and Bouvier, 2009). The SARS-CoV- 2 encoded ORF8 protein is hypothesized to downregulate the surface expression of MHC-I molecules (Zhang et al., 2020b), but we still lack a comprehensive understanding of the intrinsic capabilities of SARS-CoV-2 for immune evasion. Our study indicates that CD8+ T cells may shape the evolution of the pandemic SARS-CoV-2 virus. The majority of non-synonymous mutations found in the validated CTL escapes had not reached fixation, i.e. were present at frequencies between 0.02 and 0.42 (Fig. 1B). This could be explained by the rather short infection times of SARS-CoV-2 compared to HIV or HCV. It may also reflect on the degree to which polymorphic HLA expression affects viral spreading within the human population, as selection pressures on CTL epitopes appear to be specific for each infected individual. While it is still unclear whether these mutations may impose any fitness cost on the virus, we identified one non-synonymous mutation in the reported CTL epitope LALLLLDL that is found in an increasing number of circulating SARS-CoV-2 viruses worldwide (Hodcroft et al., 2020).

A large number of CTL epitopes for SARS-COV-2 has been described (Altmann and Boyton, 2020). Natural CTL responses in infected individuals seem to be limited to subsets of epitopes, which raises the question whether and how mutations in single epitopes affect virus control. This may be of particular importance for SARS-CoV-2 subunit vaccines that induce responses against a limited number of epitopes (Corbett et al., 2020; Liu et al., 2020; Sahin et al., 2020). In summary, our results highlight the capacity of SARS-CoV-2 to evade adaptive immune responses and provide further evidence for the impact of endogenous CTL responses and their participation in conferring protection in natural and vaccine-induced immunity.

## Acknowledgments

We thank the Biomedical Sequencing Facility at CeMM for assistance with next-generation sequencing. We thank Ursula Sinzinger, Amelie Popowitsch and Regina Sommer for excellent technical assistance; We thank Lukas Flatz, Giulio Superti-Furga, Dietmar Zehn and Rolf Zinkernagel for valuable feedback and comments.

BA was supported by the Austrian Science Fund (FWF) DK W1212. MS and AL were supported by DOC fellowships of the Austrian Academy of Sciences (No. 24813, No. 24158 and No. 24955 respectively). VG, VM and JBH received support from the Vienna Science and Technology Fund (WWTF, LS14-031). JHA was supported by the Medical-Scientific fund of the Mayor of the federal capital Vienna [grant Covid003]. C.B. and A.B. were supported by ERC Starting Grants (European Union’s Horizon 2020 research and innovation program, grant agreement numbers 679146 respectively 677006). This project was funded in part by the Vienna Science and Technology Fund (WWTF) as part of the WWTF COVID-19 Rapid Response Funding 2020 (A.B.). We declare that the research was conducted in the absence of any commercial or financial relationships that could be construed as a potential conflict of interest.

Raw BAM files were submitted for inclusion in the COVID-19 Data Portal hosted by the European Bioinformatics Institute under project number PRJEB39849. Virus sequences are deposited in the GISAID database.

## Supplementary Material

### Virus sample collection and processing

Patient samples were obtained from the Medical Universities of Vienna Institute of Virology, Medical University of Innsbruck Institute of Virology, Medical University of Innsbruck Department of Internal Medicine II and Division of Hygiene and Medical Microbiology, Central Institute for Medical-Chemical Laboratory Diagnostics Innsbruck, Klinikum Wels-Grieskirchen and the Austrian Agency for Health and Food Safety (AGES). Sample types included oropharyngeal swabs, nasopharyngeal swabs, tracheal secretion, bronchial secretion, serum and plasma. RNA was extracted using the following commercially available kits following the manufacturer’s instructions: EasyMag (bioMérieux), MagMax (Thermo Fischer), MagNA Pure LC 2.0 (Roche), AltoStar Purification Kit 1.5 (Altona Diagnostics), MagNA Pure Compact (Roche) and QIAsymphony (Qiagen). Viral RNA was reverse-transcribed with Superscript IV Reverse Transcriptase (ThermoFisher) and viral sequences were amplified with modified primer pools (Itokawa et al., 2020). PCR reactions were pooled and processed for high-throughput sequencing.

### PBMC sample collection and HLA-typing

Whole blood samples from hospitalized SARS-CoV-2-infected patients were collected at the Department of Medicine IV, Clinic Favoriten. Samples from healthy blood donors that were never exposed to SARS-CoV-2, were collected before the SARS-CoV-2 pandemic (June to November 2019). Peripheral blood mononuclear cells (PBMCs) were isolated by density gradient centrifugation and stored in liquid nitrogen until further use. HLA typing of PBMCs was carried out by Next Generation Sequencing, as described previously (Faé et al., 2019). Informed consent was obtained in accordance with the Declaration of Helsinki. The study was performed under the approval of the Ethics committee of the Medical University of Vienna, Austria (refs 2283/2019 and 1339/2017).

### Sample sequencing

AMPure XP beads (Beckman Coulter) at a 1:1 ratio were used for amplicon clean-up. Amplicon concentrations and size distribution were assessed with the Qubit Fluorometric Quantitation system (Life Technologies) and the 2100 Bioanalyzer system (Agilent), respectively. After normalization of amplicon concentrations, sequencing libraries were generated with the NEBNext Ultra II DNA Library Prep Kit for Illumina (New England Biolabs) according to manufacturer’s instructions. Library concentrations and size distribution were again assessed as indicated previously and pooled at equimolar ratios for sequencing. Sequencing was carried out on the NovaSeq 6000 platform (Illumina) on a SP flowcell with a read length of 2×250bp in paired-end mode.

### Sequencing data processing and analysis

After demultiplexing, fastq files were quality controlled using FASTQC (v. 0.11.8) (Andrews, 2010). Adapter sequences were trimmed with BBDUK from the BBtools suite (http://jgi.doe.gov/data-and-tools/bbtools). Overlaps of paired reads were corrected with the BBMERGE from BBTools. Read pairs were mapped on the combined Hg38 and SARS-CoV- 2 genome (RefSeq: NC_045512.2) using BWA-MEM with a minimal seed length of 17 (v 0.7.17) (Li and Durbin, 2009). Only reads uniquely mapping to the SARS-CoV-2 genome were retained. Primer sequences were masked with iVar (Grubaugh et al., 2019). The consensus FASTA file was generated from the binary alignment map (BAM) file using Samtools (v 1.9) (Li et al., 2009), mpileup, Bcftools (v 1.9) (Li et al., 2009), and SEQTK (https://github.com/lh3/seqtk). The read alignment file was realigned with the Viterbi method from LoFreq (v 2.1.2) for low-frequency variant calling(Wilm et al., 2012). InDel qualities were added and low-frequency variants were called with LoFreq. Variants were filtered with LoFreq and Bcftools (v 1.9) (Li, 2011). We only considered variants with a minimum coverage of 75 reads, a minimum phred-value of 90 and indels (insertions and deletions) with a HRUN of at least 4. Based on the control experiments described earlier, all analyses were performed on variants with a minimum alternative frequency of 0.02 (Popa et al., 2020). Variants were annotated with SnpEff (v 4.3) (Cingolani et al., 2012a) and SnpSift (v 4.3) (Cingolani et al., 2012b).

### Mutation Analysis

The output of LoFreq was filtered for non-synonymous-variants with a frequency cut-off of 0.02. The resulting mutations were then filtered for positions in reported CD8 T-cell epitopes. Data manipulation and plotting was carried out in R, with the packages dplyr, tidyr, ggplot2 and heatmap2.

### Identification of epitope mutations in SARS-CoV-2 genomes

Mutations in epitope regions were identified in all available protein alignment files for the SARS-CoV-2 proteins non-structural protein 3 (NSP3, n = 164819), NSP6 (n = 164806), M (n = 164846), spike (S, n = 165249), N (n = 164876) and E (n = 164847) retrieved on September 30 from the global initiative on sharing all influenza data (GISAID) database (Shu and McCauley, 2017). Protein alignment files were first filtered for protein sequences that have less than 5 % unknown amino acid positions. Epitope regions were then extracted from the alignment files and misaligned entries (>4 misaligned positions in epitope region) and protein sequences with more than 4 unknown positions in epitope region removed. Mutations in epitope regions were identified based on sequence comparison to the reference sequence “Wuhan-Hu-1” (GenBank: MN908947.3) (Wu et al., 2020).

### MHC-I binding predictions

To predict the binding strength of wildtype and mutant peptides, netMHCpan 4.1 was used (Reynisson et al., 2020). Briefly, wildtype and mutant peptide sequences were interrogated for binding to HLA-A*02:01, HLA-A*02:06 and HLA*B:40:01 with the standard settings (strong binder rank 0.5%, weak binder rank 2%). The ranks of wildtype and mutant epitopes were then compared and plotted along the heatmap of variant frequencies.

### Statistical Analysis

Statistical analysis of differences between the wildtype and mutant CD8 response was done with Wilcoxon matched-pairs signed rank test. For comparison of >1 mutant responses, a Generalized Equation Estimations (GEE) model with peptide (fixed factor) and patient (random factor) was used.

### Peptides

Peptides were purchased from JPT Peptide Technologies GmbH (Berlin, Germany) or synthesized in-house, as indicated in Table S2.

Peptides were produced in-house by solid-phase synthesis with the 9-fluorenyl-methoxy carbonyl (Fmoc)-method (CEM-Liberty, Matthews, NC, USA and Applied Biosystems, Carlsbad, CA, USA) on PEG-PS preloaded resins (Merck, Darmstadt, Germany) as previously described (Gallerano et al., 2015; Niespodziana et al., 2018) with the following alterations. After synthesis the peptides were washed with 50 ml dichloromethane (Roth, Karlsruhe, Germany), cleaved from the resins using 28.5 ml trifluoroacetic acid (Roth, Karlsruhe, Germany), 0.75 ml silane (Sigma-Aldrich, St. Louis, MO, USA) and 0.575 ml H2O for 2.5 hours at RT and precipitated into pre-chilled *tert*-butylmethylether (Merck, Darmstadt, Germany). The peptides were purified by reversed-phase high-performance liquid chromatography in a 10–70% acetonitrile gradient using a Jupiter 4 μm Proteo 90 Å, LC column (Phenomenex, Torrance, CA, USA) and an UltiMate 3000 Pump (Dionex, Sunnyvale, CA, USA) to a purity >90%. Their identities and molecular weights were verified by mass spectrometry (Microflex MALDI-TOF, Bruker, Billerica, MA, USA).

### Synthesis of HLA/Peptide complexes

cDNA encoding the extracellular domains of HLA-A*02:01 (UniProt: P01892) HLA-B*40:01 (UniProt: P01889) and beta-2-microglobulin (UniProt: P61769) were cloned without the leader sequence into pET-28b (HLA-A*02:01, HLA-B*40:01) and pHN1 (beta-2-microglobulin, β2m) for recombinant protein expression as inclusion bodies in *E. coli.* pET-28b was modified to encode a c-terminal 12x poly histidine tag (HIS12) or AviTag. Single colonies of *E. coli* (BL21) transformed with individual vectors were grown in 8l Luria-Bertani (TB) media at 37°C to an OD600 of 0.5. Protein expression was induced by addition of Isopropyl β-D- Thiogalactosid (IPTG, Sigma-Aldrich) to a final concentration of 1 μM. Cells were harvested after 4 hours of induction. Inclusion bodies containing HLA- and β2m protein were isolated, fully denatured and refolded *in vitro* in the presence of ultraviolet light-cleavable peptides (UVCP; GILGFVFJL for HLA-A*02:01; TEADVQJWL for HLA-B*40:01; J= 3-amino-3-(2- nitro)phenyl-propionic acid) to produce HLA/UVCP protein (UVCP peptides: GILGFVFJL for HLA-A*02:01, TEADVQJWL for HLA-B*40:01, J= 3-amino-3-(2-nitro)phenyl-propionic acid) (Clements et al., 2002; Garboczi et al., 1992; Toebes et al., 2006). The refolding reaction (500 ml) was dialyzed three times against 10l PBS. Dialyzed HLA/UVCP HIS12 tag proteins were purified by Ni^2+^-NTA agarose chromatography (HisTrap excel, GE Healthcare) followed by size exclusion chromatography (SEC) (Superdex 200 10/300 GL, GE Healthcare). Dialyzed HLA/UVCP AviTag proteins were concentrated to 2 ml using spin concentrators and purified by SEC. Purified HLA/UVCP AviTag proteins were biotinylated using biotin protein ligase BirA as described (Fairhead and Howarth, 2015) and further purified by SEC. The purity and integrity of all proteins was confirmed via SDS-PAGE followed by silver staining.

### UV mediated peptide exchange, Differential scanning fluorimetry (DSF) and tetramer synthesis

For peptide exchange, peptides were added to HLA/UVCP at a HLA/UVCP: peptide molar ratio of 1:20 (at a final concentration of 1.5 μM and 30 μM, respectively). For efficient cleavage, the reaction mix was placed within 5 cm from the camag^®^ uv lamp 4 (Camag) and exposed to 366 nm UV light for 2h at 4 °C followed by 16h incubation at 4 °C.

For DSF, SYPRO Orange Protein Gel Stain (Thermo Fisher Scientific, 5000x stock solution) was diluted at 4°C into the solution containing UV-treated HLA/peptide HIS12 mixtures (see above) at a final concentration of 15x SYPRO Orange Protein Gel Stain. The reaction mix was immediately transferred to pre-chilled PCR tubes and placed on a CFX 96 Real-Time PCR system (BioRad) which had been precooled to 4 °C. Samples were heated at a rate of 0.4 °C/min and relative fluorescence units (rfu) were measured every 60s in the FRET channel. Readings were plotted as negative derivative of fluorescence change vs. temperature-d(RFU)/dT.

For tetramers synthesis fluorescence-labeled streptavidin was added to UV-exchanged HLA/peptide AviTag protein solution in 10 steps as published (Altman et al., 1996). Human PBMCs were surface stained with tetramers (10 μg / ml) for 60 minutes at 4 °C followed by staining with Anti-CD8 alpha antibody (clone: OKT8) (10 μg / ml) for 30 minutes at 4 °C.

### Flow cytometry assays following 10-12d in vitro stimulation

For in vitro expansion,_cryopreserved PBMCs were thawed in pre-warmed RPMI-1640 medium (R0883, Sigma) containing 10% FBS (FBS 12-A, Capricorn), 10 mM Hepes (H0887, Sigma), 2 mM Glutamax (35050-061, Gibco), 50 IU/ml PenStrep (P4333, Sigma) and 50 IU interleukin-2 (200-02,Peprotech) at a concentration of 1×10^6^ cells/ml. PBMCs were pulsed with peptides (1 μg/ml) and cultured for 10-12d adding 100 IU interleukin-2 on day 5. In vitro expanded cells were analyzed by intracellular cytokine and cell surface marker staining. PBMCs were incubated with 2 μg/ml of peptide and 1 μg/ml anti-CD28/49d antibodies (Becton Dickinson) or with CD28/49d antibodies alone (negative control) for 6h. After 2h, 0.01 μg/ ml Brefeldin A (B7651, Sigma) was added. Staining was performed using APC/H7 anti-human CD3 (560176, BD), Pacific Blue anti-human CD4 (558116, BD), PE anti-human CD8 (555635, BD), FITC anti-human IFN-γ monoclonal antibodies (340449, BD), and Fix/Perm kit (GAS004, Invitrogen). Viable cells were determined using live/dead cell viability assay kit (L34957, Invitrogen). Cells were analyzed on a FACS Canto II cytometer (BD) and evaluated using FlowJo software v. 7.2.5 (Tree Star). The gate for detection of IFN-γ in peptide-stimulated cell samples was set in the samples with co-stimulation only.

### IFN-γ ELISpot assay

For *ex vivo* ELISpot assays, PBMCs were thawed and depleted of CD4-positive cells using magnetic microbeads coupled to anti-CD4 antibody and LD columns according to the manufacturer’s instructions (Miltenyi Biotec). A total of 1-2 x 10^5^ CD4-depleted cells per well were incubated with 2 μg/ml single peptides, AIM-V medium (negative control) or PHA (L4144, Sigma; 0,5 μg/ml; positive control) in 96-well plates coated with 1.5 μg anti-IFN-γ (3420-3-1000, Mabtech). After 45h incubation, spots were developed with 0.1 μg biotin-conjugated anti-IFN-γ (3420-6-250, Mabtech), streptavidin-coupled alkaline phosphatase (ALP; 3310-10, Mabtech, 1:1000), and 5-bromo-4-chloro-3-indolyl phosphate/nitro blue tetrazolium (BCIP/NBT; B5655, Sigma). Spots were counted using a Bio-SysBioreader 5000 Pro-S/BR177 and Bioreader software generation 10. T cell responses were considered positive when mean spot counts were at least threefold higher than the mean spot counts of three unstimulated wells.

**Fig. S1.**
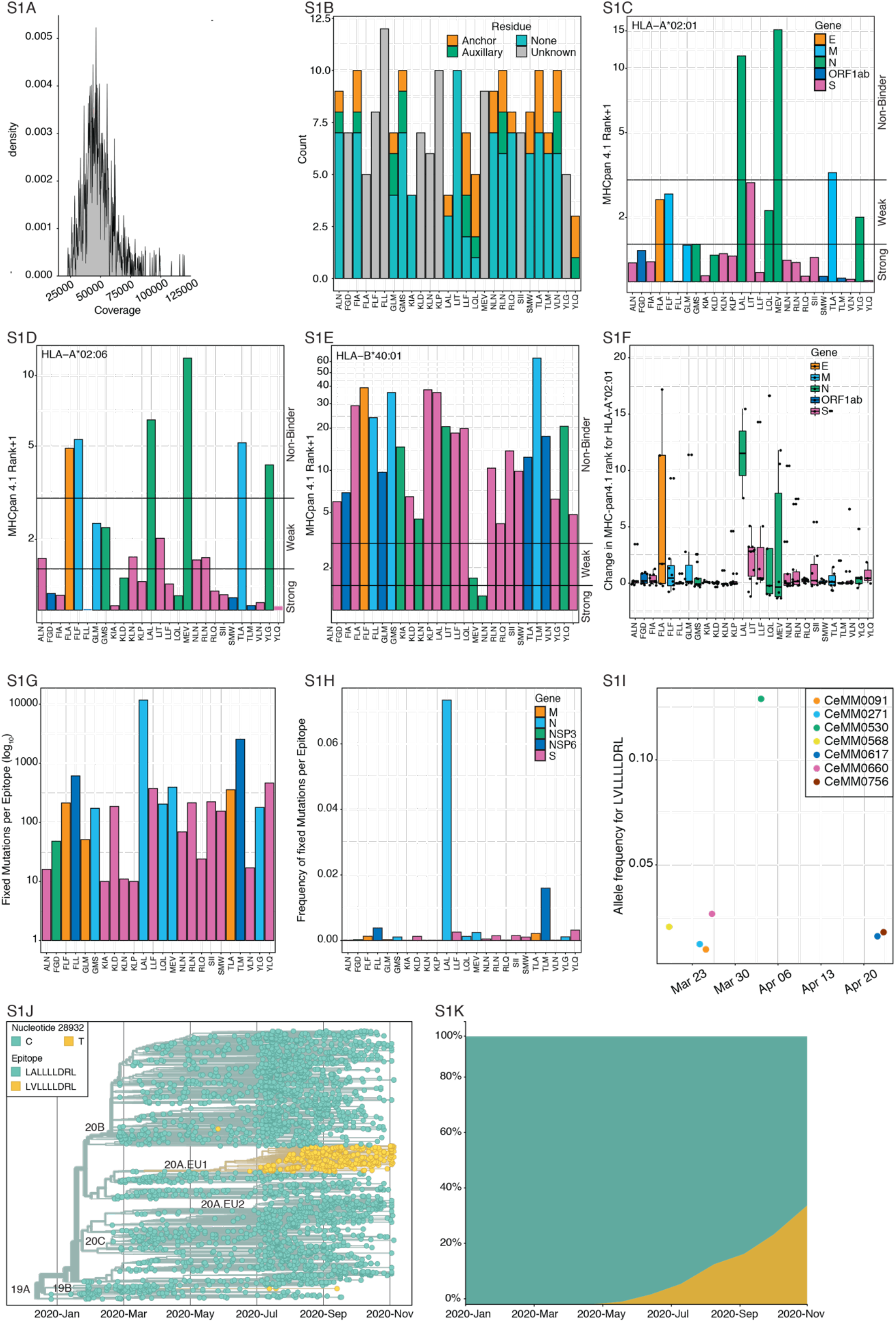
**S1A)** Coverage and read numbers of all sequenced samples. **S1B)** Mutation counts for specific residues of the epitopes. Anchor = anchor residues, auxillary = auxillary residues, none = no special residue, unknown = anchor and auxillary residues are not known for this epitope. **S1C-S1E)** netMHCpan binding predictions for the 27 wildtype epitopes to HLA-A*02:01, HLA-A*02:06 and HLA-B*40:01, respectively. **S1F)** change in netMHCpan binding predictions for the mutant wildtype epitopes to HLA-A*02:01. **S1G-S1H)** Total counts and frequency of fixed epitope mutations in the global samples. **S1I)** Allele frequency for the C28932T variant (LVLLLLDRL) in 7 patients sampled between 19.03.2020 and 23.04.2020. **S1J)** C28932T is a defining mutation of the clade A.EU1. Phylogenetic tree was generated by Nextstrain with Europe focused subsampling on 12.11.2020 **S1K)** Percentage of samples with the C28932T mutation over time.

**Fig. S2.**
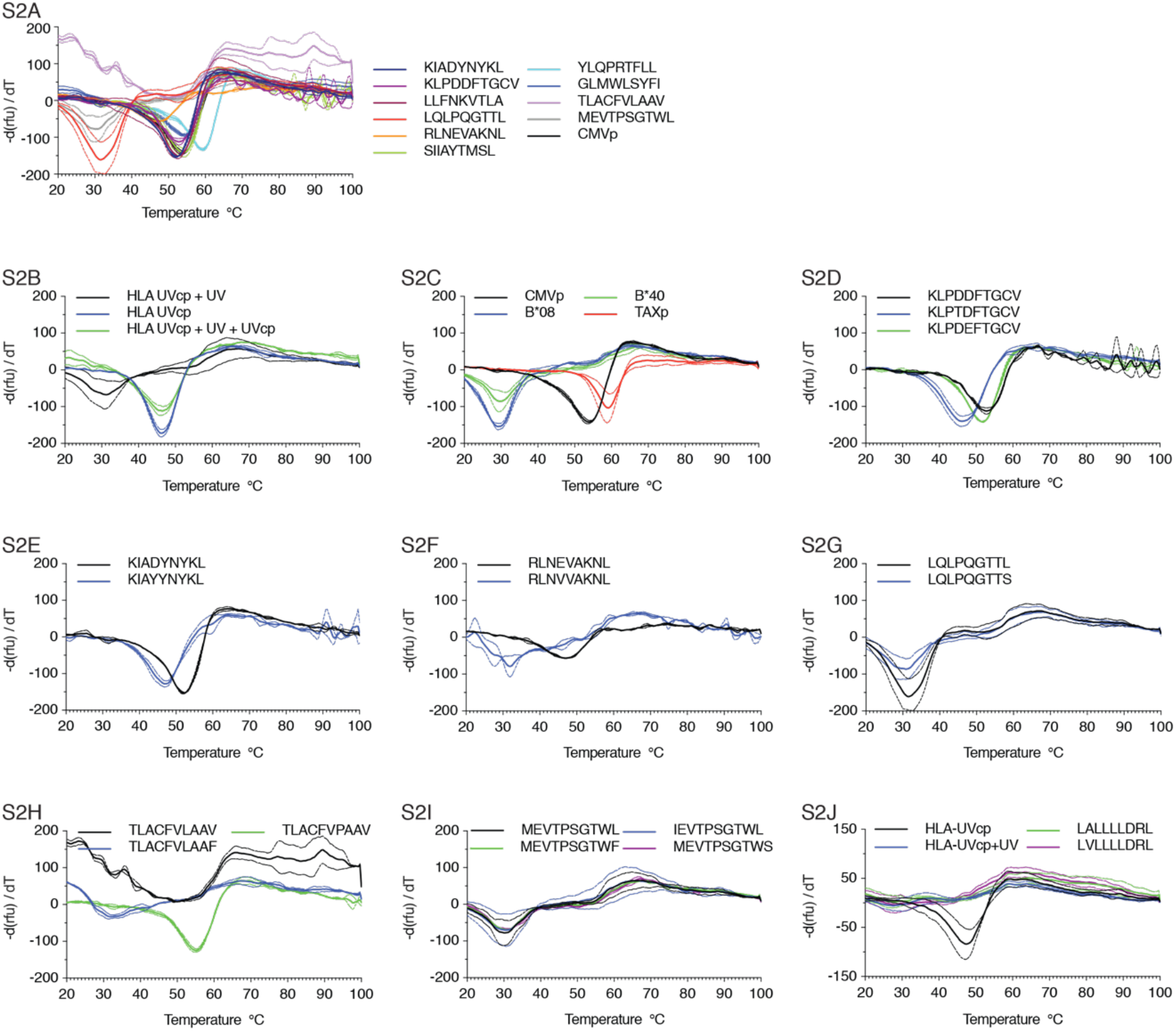
**S2A)** DSF curves for all wildtype peptides tested. **S2B)** Positive controls for the DSF assay. The black curve represents the complex of MHC and an UV cleavable peptide that was exposed to UV light, the blue curve represents the complex of MHC and an UV cleavable peptide not exposed to UV and the green curve represents the complex of MHC and an UV cleavable peptide exposed to UV light, prior to addition of another peptide. **S2C)** Additional controls for the DSF assay. Two positive controls (CMV and TAX peptides), as well as two negative controls (HLA-B epitopes) complexed with HLA-A*02:01. **S2D-S2J)** DSF data for additional mutant peptides tested. Curves for wildtype peptides are black, mutated peptides are colored. The minimal point of the curves represents the melting temperature of peptide-MHC-I complexes. **S2I)** Additional negative control for the assay. A HLA-B*40:01 epitope and its mutant forms were complexed with HLA-A*02:01

**Fig. S3.**
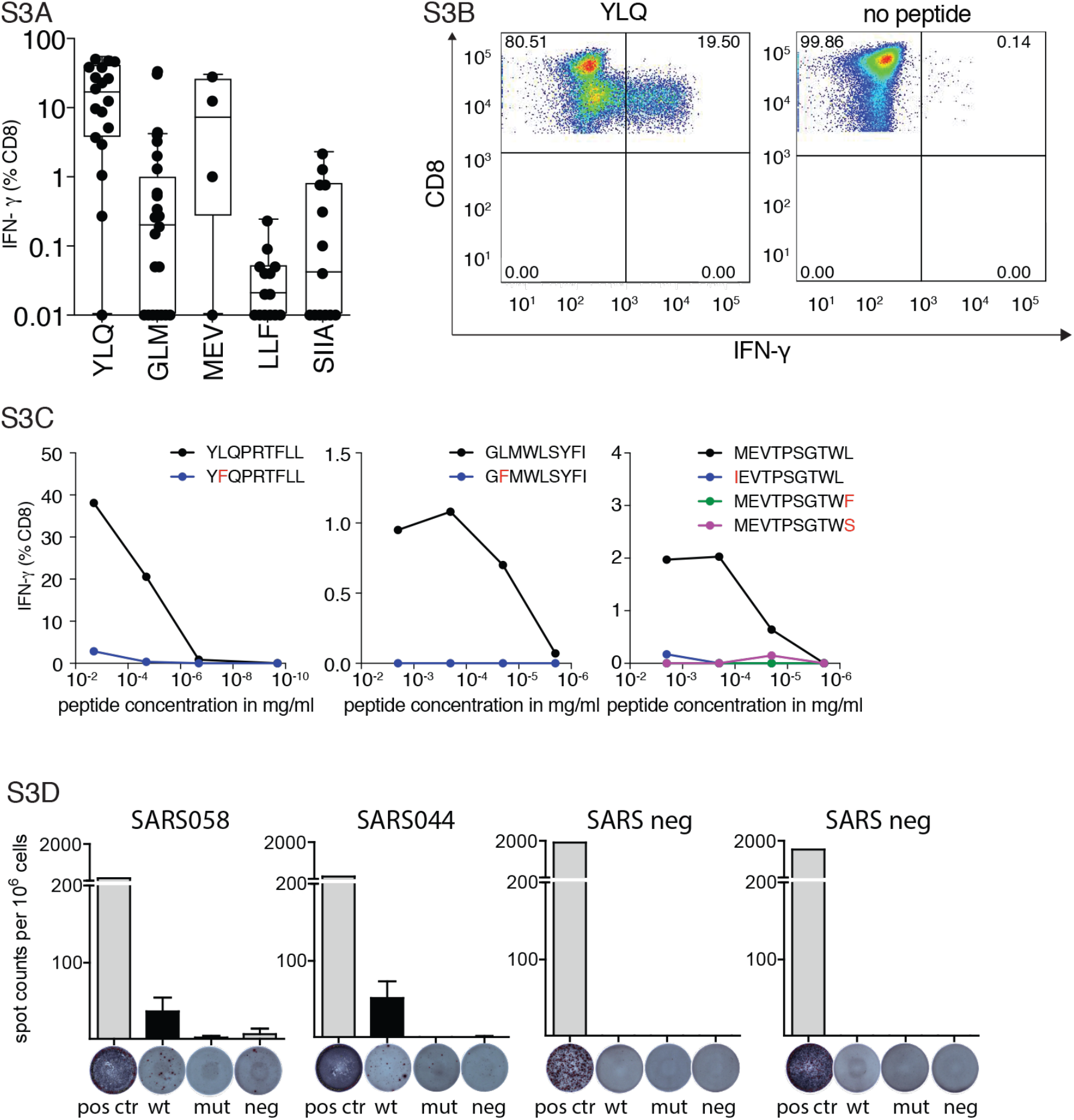
**S3A)** Intracellular cytokine staining of PBMCs after 12 days *in vitro* expansion and restimulation with wildtype peptides as indicated. **S3B)** Representative FACS plot of the ICS shown for the YLQ epitope and an unstimulated control. **S3C** Peptide titrations for three wildtype peptides and respective mutants. Peptides were tested in log_10_ and log_100_ dilutions in ICS assays. **S3D)** *Ex-vivo* ELISpot assay after restimulation with wildtype or mutant peptides in patients with detectable wildtype responses (n=6), Significance is indicated as \**P*<0.05, tested by Wilcoxon matched-pairs signed rank test **S3E)** Representative ex vivo ELISpots shown from two COVID19 patients and two healthy donors.

**Table S1.** Samples with epitope mutations at allele frequency >0.02. (separate spreadsheet)

**Table S2.**
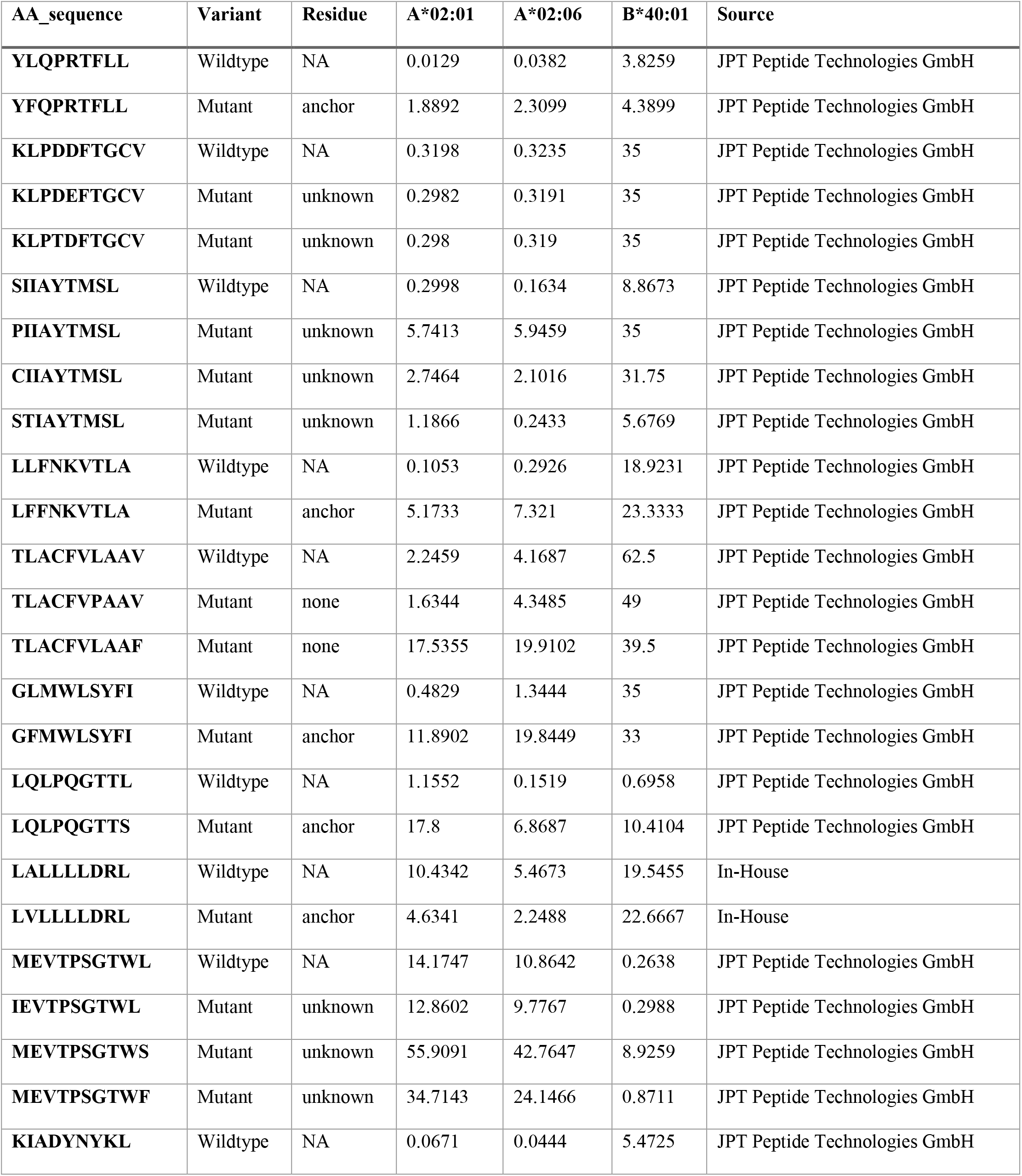
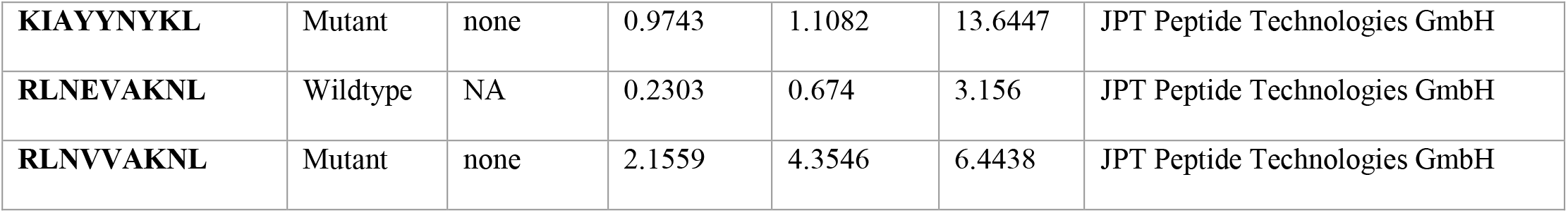
Peptides used in the study and their binding rank predicted by netMHC-pan v4.1. <0.5 strong binder, 0.5-2 weak binder, >2 non-binder.

**Table S3.** Characteristics of HLA-A*02:01/HLA-B*40:01 positive patients

|  |  |  |
| --- | --- | --- |
| Number of patients |  | **34** |
| Sex [n (%)] | Female | 11 (32) |
|  | Male | 23 (68) |
| Age [years] | Range | 22-91 |
|  | Median | 63 |
| Sample collection date | 03/2020-11/2020 |  |
| Interval symptom onset to sample collection [weeks] | Range | 1-16 |
|  | Mean | 4 |
| Chronic comorbidities [n (%)] | Hypertension | 18 (53) |
|  | Lung disease | 3 (9) |
|  | Diabetes | 14 (41) |
| Invasive ventilation [n (%)] |  | 11 (32) |
| Death [n (%)] |  | 4 (12) |

**Table S4.** HLA genotypes of included patients

| **Patient ID** | **HLA-A** | **HLA-A** | **HLA-B** | **HLA-B** | **HLA-C** | **HLA-C** |
| --- | --- | --- | --- | --- | --- | --- |
| **002** | 02:01 | 23:01 | 37:01 | 44:03 | 04:01 | 06:02 |
| **003** | 02:01 | 11:01 | 35:01 | 40:01 | 04:01 | - |
| **004** | 01:01 | 02:01 | 08:01 | 35:02 | 04:01 | 07:01 |
| **005** | 02:01 | - | 38:01 | - | 12:03 | - |
| **011** | 02:01 | - | 13:02 | 40:01 | 03:04 | 06:02 |
| **013** | 02:01 | 24:02L | 51:01 | 55:01 | 01:02 | 14:02 |
| **014** | 02:01 | 11:01 | 40:02 | 52:01 | 02:02 | 12:02 |
| **017** | 02:01 | 29:02 | 39:01 | 53:01 | 06:02 | 07:02 |
| **022** | 02:01 | 32:01 | 07:02 | 51:01 | 07:02 | 14:02 |
| **032** | 02:01 | 25:01 | 15:01 | 51:01 | 01:02 | 03:04 |
| **033** | 02:01 | 29:02 | 13:02 | 44:03 | 06:02 | 16:01 |
| **036** | 02:01 | 24:02 | 13:02 | 15:01 | 03:03 | 06:02 |
| **040** | 02:01 | 30:01 | 13:02 | 51:01 | 06:02 | 15:13 |
| **041** | 02:01 | 03:01 | 07:02 | 51:01 | 07:02 | 15:02 |
| **042** | 02:01 | 68:01 | 07:02 | 38:01 | 07:02 | 12:03 |
| **043** | 02:01 | 23:01 | 27:02 | 51:01 | 02:02 | 04:01 |
| **044** | 01:01 | 02:01 | 37:01 | 44:02 | 01:02 | 06:02 |
| **047** | 02:01 | 29:01 | 07:05 | 35:03 | 04:01 | 15:05 |
| **048** | 02:01 | 31:01 | 13:02 | 37:01 | 06:02 | - |
| **049** | 02:01 | 24:02 | 07:02 | 51:01 | 07:02 | 14:02 |
| **050** | 02:01 | 24:02 | 51:01 | 58:01 | 03:02 | 15:04 |
| **055** | 02:01 | 33:03 | 15:01 | 40:01 | 03:03 | 03:04 |
| **056** | 24:02 | 29:02 | 08:01 | 40:01 | 03:04 | 07:01 |
| **058** | 01:01 | 02:01 | 15:01 | 51:01 | 03:04 | 15:02 |
| **059** | 02:01 | 32:01 | 15:01 | 57:01 | 01:02 | 06:02 |
| **060** | 02:01 | 26:01 | 18:01 | 38:01 | 07:01 | 12:03 |
| **061** | 02:01 | 32:01 | 18:01 | 35:01 | 04:01 | 07:01 |
| **062** | 02:05 | 30:01 | 13:02 | 41:01 | 06:02 | 07:01 |
| **063** | 02:01 | 24:02 | 35:02 | 39:01 | 04:01 | 12:03 |
| **064** | 02:01 | 11:01 | 13:02 | 35:03 | 06:02 | 12:03 |
| **066** | 02:01 | 29:01 | 07:05 | 35:03 | 04:01 | 15:05 |
| **067** | 02:01 | - | 07:02 | 15:01 | 01:02 | 07:02 |
| **068** | 02:01 | - | 27:05 | 40:01 | 02:02 | 03:04 |
| **069** | 02:01 | 26:01 | 27:05 | 51:01 | 02:02 | 14:02 |
| **070\*** | 2* | *Donor HLA-typed by flow cytometry |  |  |  |  |

